# Sporulation environment drives phenotypic variation in the pathogen *Aspergillus fumigatus*

**DOI:** 10.1101/797076

**Authors:** S. Earl Kang, Michelle Momany

## Abstract

*Aspergillus fumigatus* causes more than 300,000 life-threatening infections annually and is widespread across varied environments with a single colony producing thousands of conidia, genetically-identical dormant spores. Conidia are easily wind-dispersed to new environments where they can germinate and, if inhaled by susceptible hosts, cause disease. Using high-throughput, single-cell analysis we show that germination phenotypes vary among genetically-identical individuals and that the environment of spore production determines the degree of germination heterogeneity.

---

Fungal diseases kill over 1.5 million people each year^1, 2^. Rather than spreading patient-to-patient, fungal diseases are acquired from the environment or normal flora. Nine of the ten most common agents of fungal disease can be spread via spores^2, 3^. Breaking dormancy, or germinating, is arguably the most important step in pathogenesis for these fungi. Historically studies have focused on the germination environment, addressing factors such as temperature, inoculum density, carbon source, nitrogen source, and pH^4-8^. However, despite the wide range of environments in which fungal spores are produced and their importance as disease agents, the impact of sporulation environment on germination has been largely ignored. We hypothesized that exposure to specific stresses during sporulation might lead to better germination in the same or related conditions. To test this hypothesis, we performed single-cell analysis experiments in which *A. fumigatus* was sporulated under nine environmentally- and medically-relevant conditions^9-12^ and the resulting conidia were transferred to all nine conditions for germination (Table 1). To avoid induction or selection of mutations during sporulation, we did not use serial passaging; rather, identical aliquots of inoculum were incubated for 72 h on nine types of solid medium for production of conidia and identical aliquots of conidia from each condition were transferred directly to nine types of liquid medium for germination (Supplementary Fig. 1).

**Table 1.** Sporulation and Germination Conditions.

| Abbreviation | Description | Medium | Temperature (°C) |
| --- | --- | --- | --- |
| CM | Complete medium | Nutrient-rich undefined medium containing yeast extract, glucose, nitrogen, and vitamins. | 37 |
| MM | Minimal medium | Nutrient-rich defined synthetic medium containing glucose, nitrogen and vitamins. | 37 |
| 50°C | High temperature stress | MM | 50 |
| + Cu | Copper stress | MM with 1mM CuSO<sub>4</sub> | 37 |
| + Fe | Excessive iron stress | MM with 10mM FeSO<sub>4</sub> | 37 |
| - Fe | Iron limiting stress | MM without FeSO<sub>4</sub> | 37 |
| NaCl | Osmotic or salt stress | MM with 0.5M NaCl | 37 |
| H<sub>2</sub>O<sub>2</sub> | Reactive oxygen species (ROS) stress | MM with 2mM H<sub>2</sub>O<sub>2</sub> | 37 |
| - Zn | Zinc limiting stress | MM without ZnSO<sub>4</sub> | 37 |

After 6 h incubation we used flow cytometry to detect any increase in cell size, a clear indication that germination has been initiated. The entire 9 by 9 sporulation/germination swap experiment was repeated four times. We recorded forward scatter for approximately 20,000 conidia and germlings for each condition in each replicate. For each condition, data from all replicates were concatenated and analyzed as a single population (Fig. 1, Table 2).

**Table 2.**
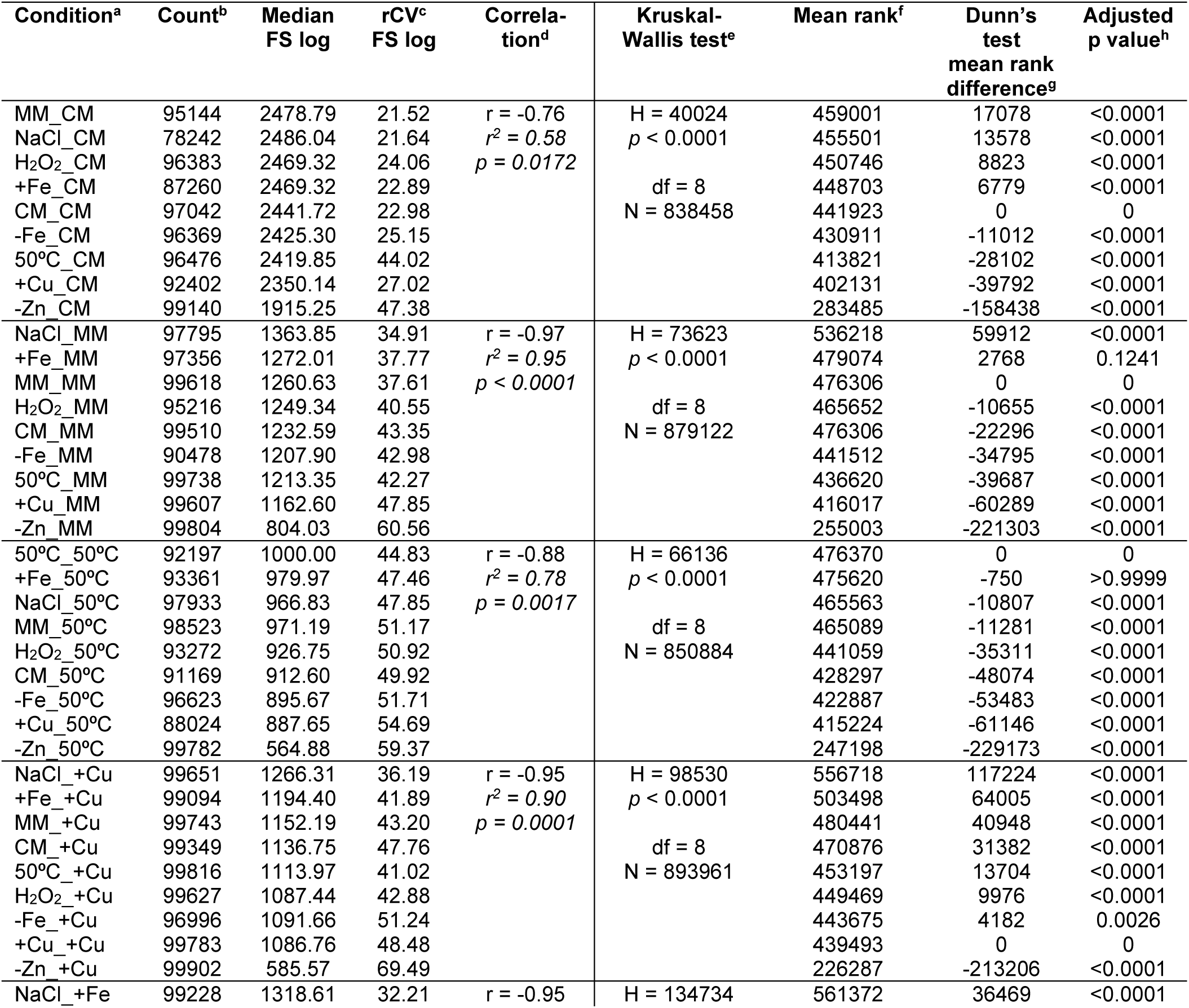
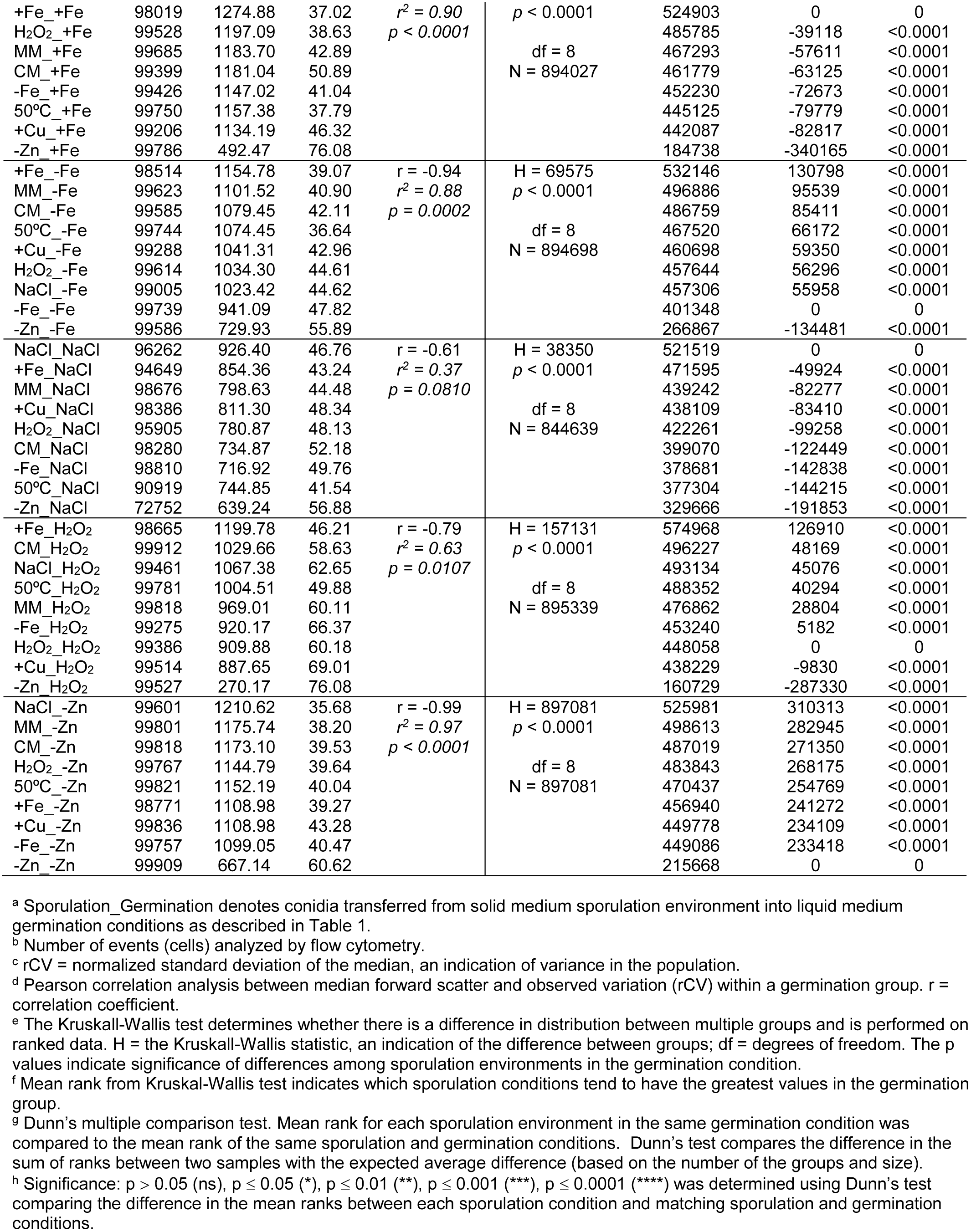
Statistical analysis of all sporulation/germination combinations grouped by germination condition.

**Figure 1.**
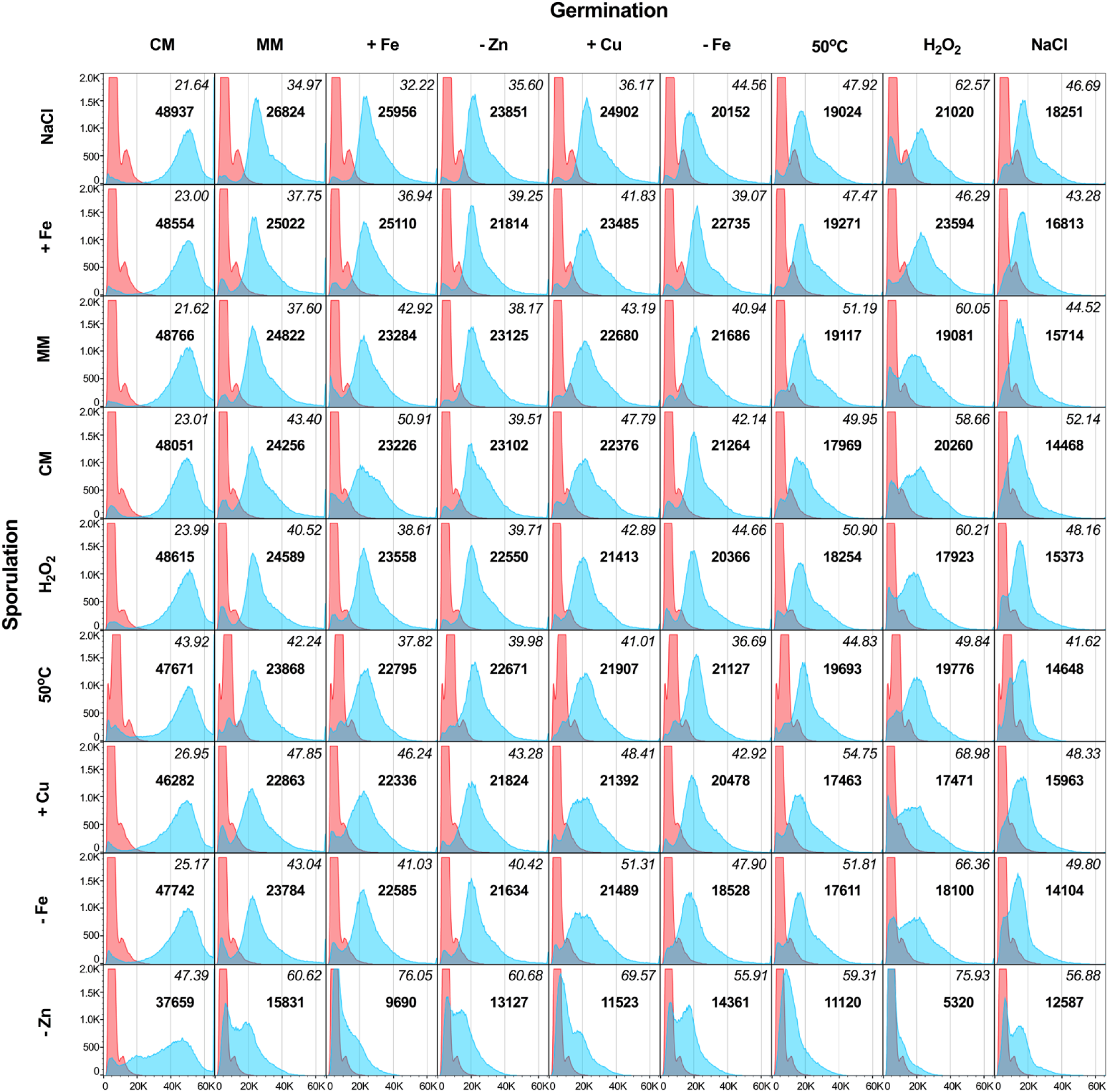
Sporulation conditions impact germination of *A. fumigatus* conidia. Conidia produced in one of nine sporulation environments were transferred to all nine conditions for germination. The X-axis shows the linear forward scatter, an indication of relative size. The Y-axis shows the number of events (cells) counted. Red peaks show forward scatter of dormant conidia from each sporulation condition measured before germination. Blue peaks show forward scatter after 6 h incubation in each germination condition. Bold values are the median of linear forward scatter values after germination. Italicized values are the robust coefficient of variation (normalized standard deviation around the median) of linear forward scatter values after germination.

Dormant conidia produced in all sporulation environments showed very similar forward scatter profiles except for conidia produced at 50°C, in which the forward scatter peak shifted slightly to the right, suggesting a larger size. Microscopic examination showed that conidia produced at 37°C were approximately 2-3 µm in diameter, while those produced at 50°C were approximately 1.5 times larger (Supplementary Fig. 2).

Not surprisingly, the rate at which conidia broke dormancy and grew varied depending on germination conditions. Conidia germinated in standard media containing sufficient metals (CM, MM) at optimal temperature (37°C) showed larger median forward scatter values than conidia germinated in media with metal limitation (−Zn, −Fe), at elevated temperature (50°C), or subjected to stressors (+Cu, +Fe, NaCl, H_2_O_2_) (Fig. 1, Table 2, and Supplementary Table 1). Across all sporulation environments, conidia broke dormancy and grew more quickly in CM germination medium than in any other germination condition. Conidia germinated in 0.5M NaCl (osmotic stress) generally broke dormancy and grew more slowly than those in other germination conditions. These results are consistent with previous work showing that rich medium and non-stressful conditions during germination favor more rapid dormancy breaking and growth^13,14,15^.

In addition to the expected contribution of germination conditions, the rate at which conidia broke dormancy and grew varied depending on sporulation environment. The sporulation environments that favored rapid dormancy breaking and growth were not the same as the germination conditions that favored it. As discussed above, 0.5M NaCl during germination resulted in reduced dormancy breaking and growth. In contrast, osmotic stress imposed by 0.5M NaCl during sporulation resulted in conidia that broke dormancy and grew more quickly across germination conditions. In addition to NaCl medium, sporulation on MM or +Fe medium generally improved dormancy breaking and growth when compared to conidia from all other sporulation environments. Conidia from +Cu, −Fe, and −Zn sporulation environments generally performed significantly worse when compared to conidia from MM condition (Supplementary Table 2) suggesting that proper metal homeostasis is necessary during sporulation as well as germination. These results show for the first time that the sporulation environment impacts the ability of a medically-important fungus to break dormancy and grow across multiple germination environments.

While we predicted that forward scatter peaks might shift left or right with changes in germination or sporulation conditions, we were surprised to see striking differences in the widths and shapes of peaks depending on sporulation environment. *A. fumigatus* conidia are clonal, with each conidium in a colony containing a single genetically-identical nucleus produced by mitosis. Previous work has shown that conidia remain dormant until they are exposed to a carbon source and water^6^, at which time individuals in the population synchronously break dormancy and start growth, with rough synchrony maintained through at least the first 12 hours^16^. Thus, we expected that individual conidia produced in the same sporulation environment would break dormancy and grow synchronously, giving rise to relatively narrow peaks. The observed wide peaks show that genetically-identical conidia within the same population break dormancy and grow at different rates. The dramatic leftward shift of post-germination peaks for sporulation conditions such as −Zn medium could be explained if Zn deficiency during sporulation killed conidia. However, viability assays with fluorescein diacetate and propidium iodide showed that conidia sporulated on MM and on −Zn media contained very similar, low numbers of propidium iodide stained cells and that most of the conidia that did not enlarge during germination were not dead (Supplementary Table 3).

To better understand the range of individual variation within genetically-identical clonal populations of conidia, we compared the robust coefficient of variation (rCV, the normalized standard deviation of the median) for forward scatter of each sporulation/germination pair (Table 2 and Supplementary Table 2). Conidia that were produced on NaCl, +Fe, 50°C, and MM sporulation media showed lower rCV values and narrower forward scatter peaks across germination conditions, indicating less variation among individuals in those populations. Conidia from −Zn, −Fe, CM, +Cu, and H2O2 sporulation medium showed higher rCV values and wider forward scatter peaks across germination conditions, indicating more variation among individuals in those populations (Fig. 1, Supplementary Table 2). Taken together with median forward scatter values this shows that conidia that germinate faster tend to germinate more synchronously. Indeed, there was a negative correlation between median growth and variation in growth across most conditions (Table 2). The correlation between sporulation medium and variation was much stronger than the correlation between germination medium and variation (Supplementary Table 1, Supplementary Table 2) consistent with the idea that the environment of sporulation drives germination variation.

Our results show for the first time that the environment of spore production impacts the germination of *A. fumigatus* conidia and that genetically-identical conidia within a population vary in the rate of breaking dormancy and growth. That genetically-identical individuals show phenotypic variation that is increased by environmental stress suggests *A. fumigatus* might employ a bet-hedging strategy to ensure survival of progeny in varied hostile environments, including the lungs of susceptible human hosts. Previous work showed that the surface layer of dormant *A. fumigatus* conidia mask recognition by the host immune system. It is only when dormancy is broken and germination occurs that this surface layer is breached and host defenses are activated^17^. Other studies in immunosuppressed mice showed that an *A. fumigatus* isolate with slower germination survived in macrophages and was more virulent than an isolate with faster germination^18,19^. A bet-hedging strategy built on variation in germination rate could allow slow germinators within a population of *A. fumigatus* conidia to avoid the host immune system and initiate infection. It seems likely that this bet-hedging strategy would also be used by the many other fungal pathogens that produce large quantities of wind-dispersed spores.

**Supplementary Table 1.** Statistical analysis of germination conditions.

| Germination<sup>a</sup> | Counts<sup>b</sup> | Median<br>FS log | rCV<sup>c</sup><br>FS log | Corre-<br>lation<sup>d</sup> | Kruskal-Wallis test<sup>e</sup> | Mean rank<sup>f</sup> | Dunn's<br>test<br>mean rank<br>difference<sup>g</sup> | Adjusted<br>p value<sup>h</sup> |
| --- | --- | --- | --- | --- | --- | --- | --- | --- |
| _CM | 838458 | 2404 | 28.58 | $r = -0.77$<br>$r^2 = 0.59$<br>$p = 0.0159$ | H = 1578766<br>df = 8<br>$p < 0.0001$<br>N = 7888209 | 6496947 | 2135891 | <0.0001 |
| _MM | 879122 | 1213 | 45.10 |  |  | 4361056 | 0 | 0 |
| _<sup>+</sup>Fe | 894027 | 1152 | 48.64 |  |  | 4049783 | -311274 | <0.0001 |
| _<sup>-</sup>Zn | 897081 | 1106 | 43.18 |  |  | 4027523 | -333533 | <0.0001 |
| _<sup>+</sup>Cu | 893961 | 1102 | 48.76 |  |  | 3926138 | -434873 | <0.0001 |
| _<sup>-</sup>Fe | 894698 | 1030 | 44.98 |  |  | 3705829 | -655228 | <0.0001 |
| _50°C | 850884 | 907.8 | 53.52 |  |  | 3205174 | -1155883 | <0.0001 |
| _H2O2 | 895339 | 926.4 | 67.61 |  |  | 3171735 | -1189321 | <0.0001 |
| _NaCl | 844639 | 784.4 | 48.86 |  |  | 2610009 | -1751048 | <0.0001 |

<sup>a</sup> \_Germination denotes concatenated data of all conidia transferred from each of the nine solid medium sporulation environments into the designated liquid medium germination condition as described in Table 1.
<sup>b</sup> Number of events (cells) analyzed by flow cytometry.
<sup>c</sup> rCV = normalized standard deviation of the median, an indication of variance in the population.
<sup>d</sup> Pearson correlation analysis between median forward scatter and observed variation (rCV) between germination groups. $r$ = correlation coefficient.
<sup>e</sup> The Kruskal-Wallis test determines whether there is a difference in distribution between multiple groups and is performed on ranked data. $H$ = the Kruskal-Wallis statistic, an indication of the difference between groups; $df$ = degrees of freedom. The $p$ values indicate significance of differences between germination conditions.
<sup>f</sup> Mean rank from Kruskal-Wallis test indicates which germination conditions tend to have the greatest values.
<sup>g</sup> Dunn's multiple comparison test. Mean rank for each germination condition was compared to the mean rank of germination in MM (the base medium for all conditions). Dunn's test compares the difference in the sum of ranks between two samples with the expected average difference (based on the number of the groups and size).
<sup>h</sup> Significance: $p > 0.05$ (ns), $p \leq 0.05$ (\*), $p \leq 0.01$ (\*\*), $p \leq 0.001$ (\*\*\*), $p \leq 0.0001$ (\*\*\*\*) was determined using Dunn's test comparing the difference in the mean ranks between each germination condition and MM (the base medium).

**Supplementary Table 2.**
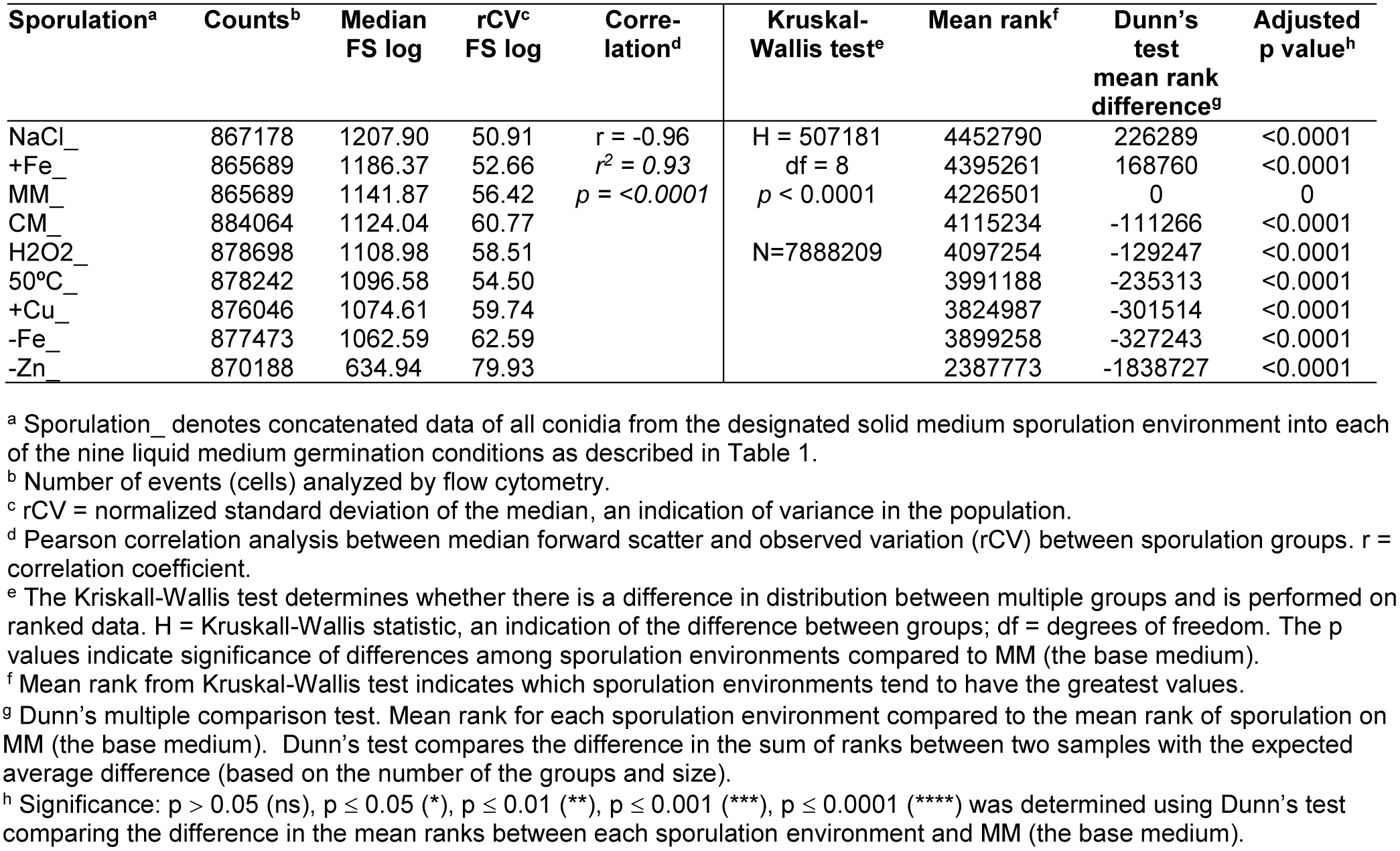
Statistical analysis of sporulation environments.

| Sporulation<sup>a</sup> | Counts<sup>b</sup> | Median<br>FS log | rCV<sup>c</sup><br>FS log | Corre-<br>lation<sup>d</sup> | Kruskal-Wallis test<sup>e</sup> | Mean rank<sup>f</sup> | Dunn's<br>test<br>mean rank<br>difference<sup>g</sup> | Adjusted<br>p value<sup>h</sup> |
| --- | --- | --- | --- | --- | --- | --- | --- | --- |
| NaCl_ | 867178 | 1207.90 | 50.91 | $r = -0.96$<br>$r^2 = 0.93$<br>$p < 0.0001$ | H = 507181<br>df = 8<br>$p < 0.0001$<br>N=7888209 | 4452790 | 226289 | <0.0001 |
| +Fe_ | 865689 | 1186.37 | 52.66 |  |  | 4395261 | 168760 | <0.0001 |
| MM_ | 865689 | 1141.87 | 56.42 |  |  | 4226501 | 0 | 0 |
| CM_ | 884064 | 1124.04 | 60.77 |  |  | 4115234 | -111266 | <0.0001 |
| H2O2_ | 878698 | 1108.98 | 58.51 |  |  | 4097254 | -129247 | <0.0001 |
| 50°C_ | 878242 | 1096.58 | 54.50 |  |  | 3991188 | -235313 | <0.0001 |
| +Cu_ | 876046 | 1074.61 | 59.74 |  |  | 3824987 | -301514 | <0.0001 |
| -Fe_ | 877473 | 1062.59 | 62.59 |  |  | 3899258 | -327243 | <0.0001 |
| -Zn_ | 870188 | 634.94 | 79.93 |  |  | 2387773 | -1838727 | <0.0001 |

<sup>a</sup> Sporulation\_ denotes concatenated data of all conidia from the designated solid medium sporulation environment into each of the nine liquid medium germination conditions as described in Table 1.
<sup>b</sup> Number of events (cells) analyzed by flow cytometry.
<sup>c</sup> rCV = normalized standard deviation of the median, an indication of variance in the population.
<sup>d</sup> Pearson correlation analysis between median forward scatter and observed variation (rCV) between sporulation groups. $r$ = correlation coefficient.
<sup>e</sup> The Kruskal-Wallis test determines whether there is a difference in distribution between multiple groups and is performed on ranked data. $H$ = Kruskal-Wallis statistic, an indication of the difference between groups; $df$ = degrees of freedom. The $p$ values indicate significance of differences among sporulation environments compared to MM (the base medium).
<sup>f</sup> Mean rank from Kruskal-Wallis test indicates which sporulation environments tend to have the greatest values.
<sup>g</sup> Dunn's multiple comparison test. Mean rank for each sporulation environment compared to the mean rank of sporulation on MM (the base medium). Dunn's test compares the difference in the sum of ranks between two samples with the expected average difference (based on the number of the groups and size).
<sup>h</sup> Significance: $p > 0.05$ (ns), $p \leq 0.05$ (\*), $p \leq 0.01$ (\*\*), $p \leq 0.001$ (\*\*\*), $p \leq 0.0001$ (\*\*\*\*) was determined using Dunn's test comparing the difference in the mean ranks between each sporulation environment and MM (the base medium).

**Supplementary Table 3.**
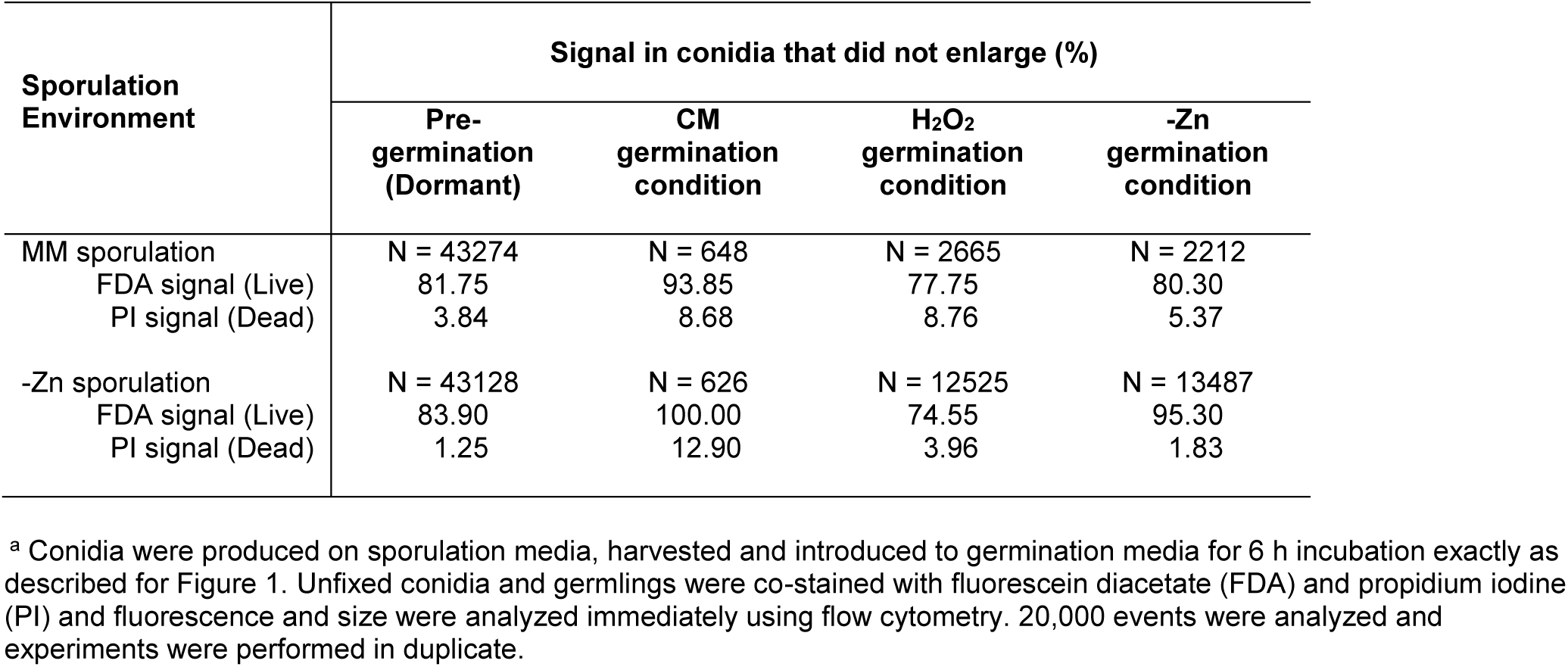
Viability Assay^a^.

| Sporulation Environment | Signal in conidia that did not enlarge (%) |  |  |  |
| --- | --- | --- | --- | --- |
|  | Pre-germination (Dormant) | CM germination condition | H<sub>2</sub>O<sub>2</sub> germination condition | -Zn germination condition |
| MM sporulation | N = 43274 | N = 648 | N = 2665 | N = 2212 |
| FDA signal (Live) | 81.75 | 93.85 | 77.75 | 80.30 |
| PI signal (Dead) | 3.84 | 8.68 | 8.76 | 5.37 |
| -Zn sporulation | N = 43128 | N = 626 | N = 12525 | N = 13487 |
| FDA signal (Live) | 83.90 | 100.00 | 74.55 | 95.30 |
| PI signal (Dead) | 1.25 | 12.90 | 3.96 | 1.83 |

<sup>a</sup> Conidia were produced on sporulation media, harvested and introduced to germination media for 6 h incubation exactly as described for Figure 1. Unfixed conidia and germlings were co-stained with fluorescein diacetate (FDA) and propidium iodine (PI) and fluorescence and size were analyzed immediately using flow cytometry. 20,000 events were analyzed and experiments were performed in duplicate.

**Supplementary Figure 1.**
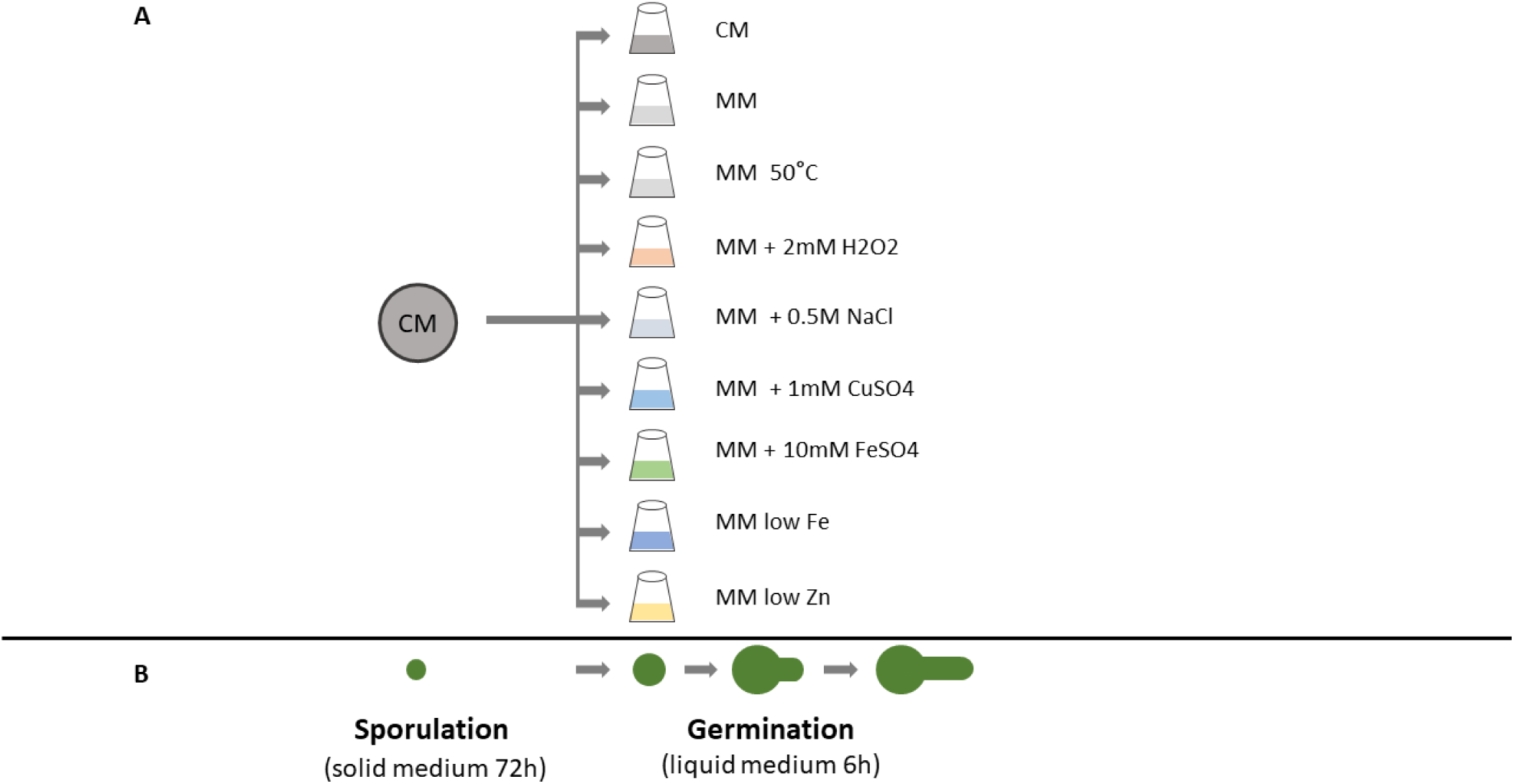
Sporulation/Germination swap assay. (A) Conidia isolated from a single sporulation condition on solid medium (CM, indicted by circle) were aliquoted into all germination conditions in liquid medium (indicated by flask shapes). The same process was repeated with conidia from each of the nine sporulation conditions being transferred to all nine germination conditions. Different colors represent different sporulation or germination conditions as indicated. (B) Diagram of relative conidium size and shape after sporulation, during germination, and for the first 6 h of growth. Dormant conidia are 2-3 microns in diameter. Upon exposure to carbon and water they break dormancy and begin to increase in size with swelling and germ tube emergence.

**Supplementary Figure 2.**
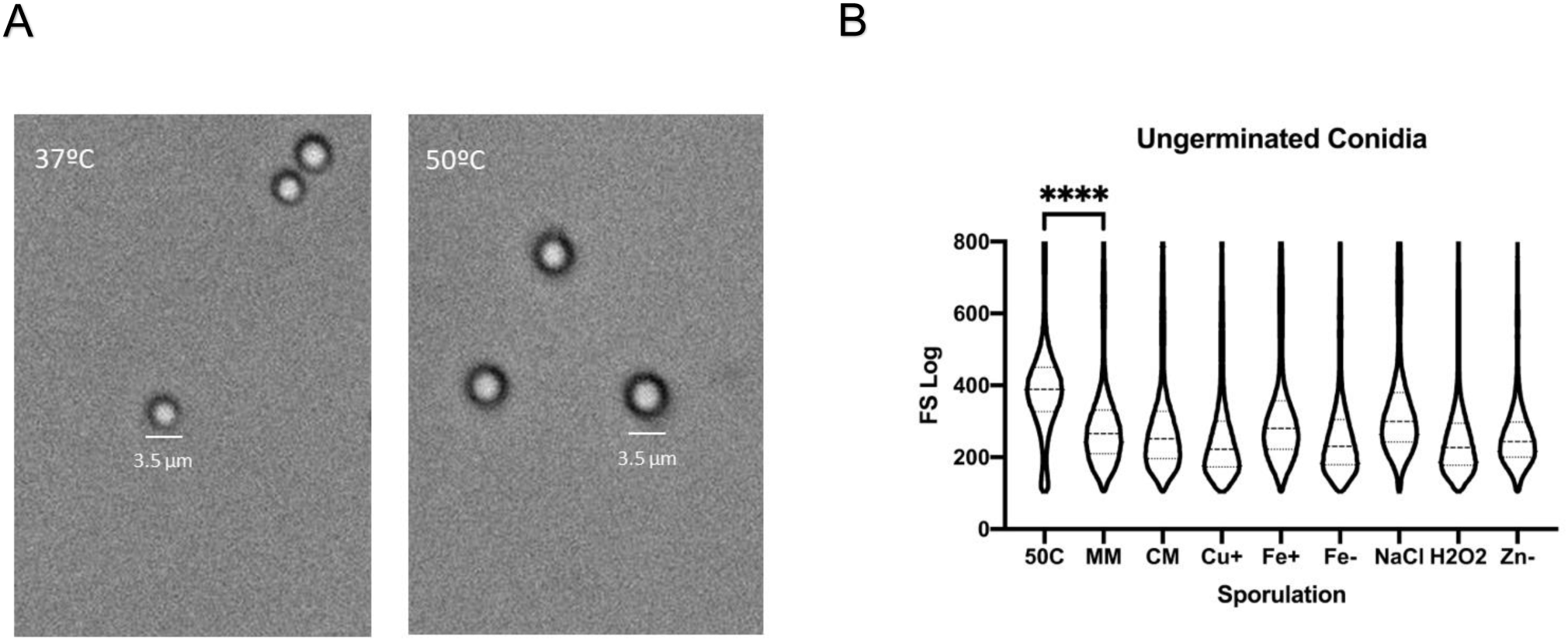
Conidia produced at 50°C are larger. (A) Light microscopy of conidia sporulated at 37**°**C and 50**°**C on minimal medium, 1,000X magnification. (B) Violin plot of forward scatter log scaled values of dormant (ungerminated) conidia from all sporulation environments. Dashed line represents median. Dotted line represents quantile at 25% and 75%. Kruskal-Wallis test followed by one-sided Dunn’s multiple comparison tests. Significance: p ≤ 0.0001 (****)

## Methods

### Fungal Strains, cultivation and preparation of conidia

*Aspergillus fumigatus* CEA10 was cultivated on 1.5% agar solid complete media (CM) or minimal media (MM) as previously described^20^with modifications as described in Table 1. For conidial stock preparation, conidia were produced on complete media, harvested in sterile water, and 1×10^6^ conidia in 500 µl of ddH2O was plated in a homogenous layer on 25ml of solid 1% glucose *Aspergillus* minimal media with modifications described in Table 1 in 90mm plates in 3 technical replicates. Plates were incubated in the dark, stored upside down at 37°C or 50°C for 72hrs. *A. fumigatus* conidia from 3 plates were harvested by overlaying plates with 25ml sterile ddH2O, combining conidia and filtering through 22-25 µm Miracloth (MilliporeSigma, St. Louis, MO, USA). Conidia were washed twice in ddH2O and counted using a hemocytometer.

### Germination assay

Conidia from 3 plates were pooled and identical aliquots of 3-5 × 10^5^ C/ml were added to liquid germination conditions described in Table 1^7^. Cultures were incubated for 6hrs at 37C or 50°C @ 250 rpm in dark, then fixed with 2.5% formaldehyde. 81 conditions were analyzed in total. Controls included conidia fixed at 0hr in liquid germination conditions.

### Analysis of germination / Flow cytometry

Flow cytometry was performed at the Center for Tropical and Emerging Global Diseases Cytometry Shared Resource Laboratory at the University of Georgia on a CyAn ADP using Summit, version 4.3 (Beckman Coulter, Fullerton, CA, USA). Between 20,000 – 25,0000 events (cells) were analyzed in four replicates for each fixed pre- and post-germination sample. Due to the sensitivity of flow cytometry and small particulates in the germinated samples, forward scatter and side scatter values smaller than fixed ungerminated conidia were filtered from the analysis. FlowJo flow cytometry analysis software, version 10 (Tree Star, Ashland, OR, USA) was used for analysis and histogram. Histogram represents the linear scaled forward scatter data to better visualize the variation in germination. Morphologies were verified using Amnis ImageStream (Amnis MerckMillipore Sigma, Seattle, WA, USA).

### Viability assay - Live / dead staining

For viability assays, two replicates of unfixed cells (conidia and germlings) were co-stained with 10 ug/ml fluorescein diacetate (FDA) and 2 ug/ml propidium iodine (PI) for 5 minutes in the dark, then 20,000 events were analyzed immediately using flow cytometry to measure size (forward scatter) and fluorescence. Controls included unstained and fluorescein diacetate (FDA), propidium iodine (PI), and FDA+PI stained live and dead (ethanol-killed) cells.

### Statistical analysis

Forward scatter scaled linear or log data was combined for each condition from all replicates. Linear and log data were checked for normality using D’Agostino-Pearson test ^21^. Due to nonparametric distribution, comparison between multiple groups were analyzed by Kruskal-Wallis test followed by one-sided Dunn’s multiple comparison test^22^ using GraphPad Prism version 8 (GraphPad Software, La Jolla, CA, USA). Robust coefficient of variance (rCV) was calculated using 100 * 1/2 (Intensity [at 84.13 percentile] – Intensity [at 15.87 percentile]) / Median using FlowJo v10 (Tree Star, Ashland, OR, USA). Pearson correlation analysis followed by a two-tailed test was performed to assess the relationship between median log forward scatter (growth) and rCV (variation) in a given germination condition using GraphPad Prism version 8 (GraphPad Software, La Jolla, CA, USA).

## Acknowledgments

We thank Julie Nelson at the UGA CTEGD Cytometry Shared Resource Laboratory for assistance with flow analysis, Douda Bensasson (UGA) for assistance with statistical analysis, and the UGA Department of Plant Biology and Franklin College of Arts and Sciences for funding.

## References

1. Bongomin, F., Gago, S., Oladele, R.O. & Denning, D.W. Global and Multi-National Prevalence of Fungal Diseases-Estimate Precision. J Fungi (Basel) 3, (2017).

2. Brown, G.D., et al. Hidden killers: human fungal infections. Sci Transl Med 4, 165rv13 (2012).

3. Botts, M.R. & Hull, C.M. Dueling in the lung: how Cryptococcus spores race the host for survival. Curr Opin Microbiol 13, 437–42 (2010).

4. Brown, W. On the germination and growth of fungi at various temperatures and in various concentrations of oxygen and of carbon dioxide. Ann Bot-London 36, 257–283 (1922).

5. Loo, M. Some required events in conidial germination of Neurospora crassa. Dev Biol 54, 201–13 (1976).

6. Osherov, N. & May, G.S. The molecular mechanisms of conidial germination. Fems Microbiol Lett 199, 153–60 (2001).

7. Araujo, R. & Rodrigues, A.G. Variability of germinative potential among pathogenic species of Aspergillus. J Clin Microbiol 42, 4335–7 (2004).

8. Wang, Z., et al. Metabolism and Development during Conidial Germination in Response to a Carbon-Nitrogen-Rich Synthetic or a Natural Source of Nutrition in Neurospora crassa. MBio 10, (2019).

9. Errasquin, E.L., Patino, B., Fernandez, R.M. & Vazquez, C. Occurrence of Aspergillus fumigatus in a compost polluted with heavy metals. Microbiology of Composting 487–494 (2002).

10. Tepsic, K., Gunde-Cimerman, N. & Frisvad, J.C. Growth and mycotoxin production by Aspergillus fumigatus strains isolated from a saltern. Fems Microbiol Lett 157, 9–12 (1997).

11. Haas, H. Iron - A Key Nexus in the Virulence of Aspergillus fumigatus. Front Microbiol 3, 28 (2012).

12. Amich, J. & Calera, J.A. Zinc acquisition: a key aspect in Aspergillus fumigatus virulence. Mycopathologia 178, 379–85 (2014).

13. Schmit JC B.S. Biochemical genetics of Neurospora crassa conidial germination. Bacteriological Reviews 40, 1–41 (1976).

14. Meletiadis, J., Meis, J.F., Mouton, J.W., Verweij, P.E. Analysis of growth characteristics of filamentous fungi in different nutrient media. Journal of Clinical Microbiology 39, (2001).

15. Osherov, N., Conidial Germination in Aspergillus fumigatus, in Aspergillus Fumigatus and Aspergillosis (eds Latgé, J.P., Steinbach, W.J.) 131–142 (ASM Press: Washington, D.C., 2009).

16. Momany, M. & Taylor, I. Landmarks in the early duplication cycles of Aspergillus fumigatus and Aspergillus nidulans: polarity, germ tube emergence and septation. Microbiology 146 Pt 12, 3279–84 (2000).

17. Aimanianda, V., et al. Surface hydrophobin prevents immune recognition of airborne fungal spores. Nature 460, 1117–21 (2009).

18. Amarsaikhan, N., et al. Isolate-dependent growth, virulence, and cell wall composition in the human pathogen Aspergillus fumigatus. PLoS One 9, e100430 (2014).

19. Rosowski, E.E., et al. Macrophages inhibit Aspergillus fumigatus germination and neutrophil-mediated fungal killing. PLoS Pathog 14, e1007229 (2018).

20. Momany, M., Westfall, P.J. & Abramowsky, G. Aspergillus nidulans swo mutants show defects in polarity establishment, polarity maintenance and hyphal morphogenesis. Genetics 151, 557–567 (1999).

21. Dagostino, R.B., Belanger, A. & Dagostino, R.B. A Suggestion for Using Powerful and Informative Tests of Normality. American Statistician 44, 316–321 (1990).

22. Dunn, O.J. Multiple Comparisons Using Rank Sums. Technometrics 6, 241–252 (1964).

